# The genetic basis of sex determination in grapevines (*Vitis spp.*)

**DOI:** 10.1101/2019.12.11.861377

**Authors:** Mélanie Massonnet, Noé Cochetel, Andrea Minio, Amanda M. Vondras, Aline Muyle, Jerry Lin, Jadran F. Garcia, Yongfeng Zhou, Massimo Delledonne, Summaira Riaz, Rosa Figueroa-Balderas, Brandon S. Gaut, Dario Cantu

## Abstract

Sex determination in grapevine evolved through a complex succession of switches in sexual systems. Phased genomes built with single molecule real-time sequencing reads were assembled for eleven accessions of cultivated hermaphrodite grapevines and dioecious males and females, including the ancestor of domesticated grapevine and other related wild species. By comparing the phased sex haplotypes, we defined the sex locus of the *Vitis* genus and identified polymorphisms spanning regulatory and coding sequences that are in perfect association with each sex-type throughout the genus. These findings identified a novel male-fertility candidate gene, *INP1*, and significantly refined the model of sex determination in *Vitis* and its evolution.

## Introduction

Plants display a great variety of sexual systems (Barrett, 2002). Though common in animals, strictly male or female individuals (dioecy) are rare in flowering plants. Dioecy is phylogenetically widespread (Käfer *et al*., 2017), but occurs in only about 5% of angiosperms, whereas monoecious individuals carrying male and female flowers and plants with hermaphrodite flowers are more common (Renner, 2014). In angiosperms, non-recombining genetic sex loci determine sex (Charlesworth and Charlesworth, 1978; Charlesworth, 2016). However, the genes involved in sex determination are largely unknown. Evidence to-date supports several models, including a two-locus model (Westergaard, 1958) in *Fragaria virginiana* (Spigler *et al.*, 2008), *Silene latifolia* (Fujita *et al.*, 2011) and *Actinidia* spp. (Akagi *et al.*, 2019), a single switch gene in *Diospyros lotus* (Akagi *et al.*, 2014) and others (Renner, 2016; Pannell, 2017; Henry *et al*., 2018).

Three flower sexes exist in grapevines (*Vitis* spp.; Negi and Olmo, 1970; **Fig. 1a, b**). There are (i) male flowers, which are sterile females with pistils reduced to a small ovary and neither stigma nor style development, (ii) female flowers, with reflexed anthers and stamens releasing few sterile pollen grains (Gallardo *et al.*, 2009), and (iii) hermaphrodite “perfect” flowers, whose pistils are functional and the erect stamens bear fertile pollen grains. Interestingly, cultivated grapevines (*Vitis vinifera* ssp. *vinifera*; hereafter *Vv vinifera*) are hermaphrodite (Pratt, 1971), a trait selected during the domestication of its dioecious wild ancestor, *Vitis vinifera* ssp. *sylvestris* (hereafter *Vv sylvestris*), nearly 8,000 years ago (This *et al.*, 2006; McGovern *et al.*, 2017; Zhou *et al.*, 2019a).

**Fig. 1:**
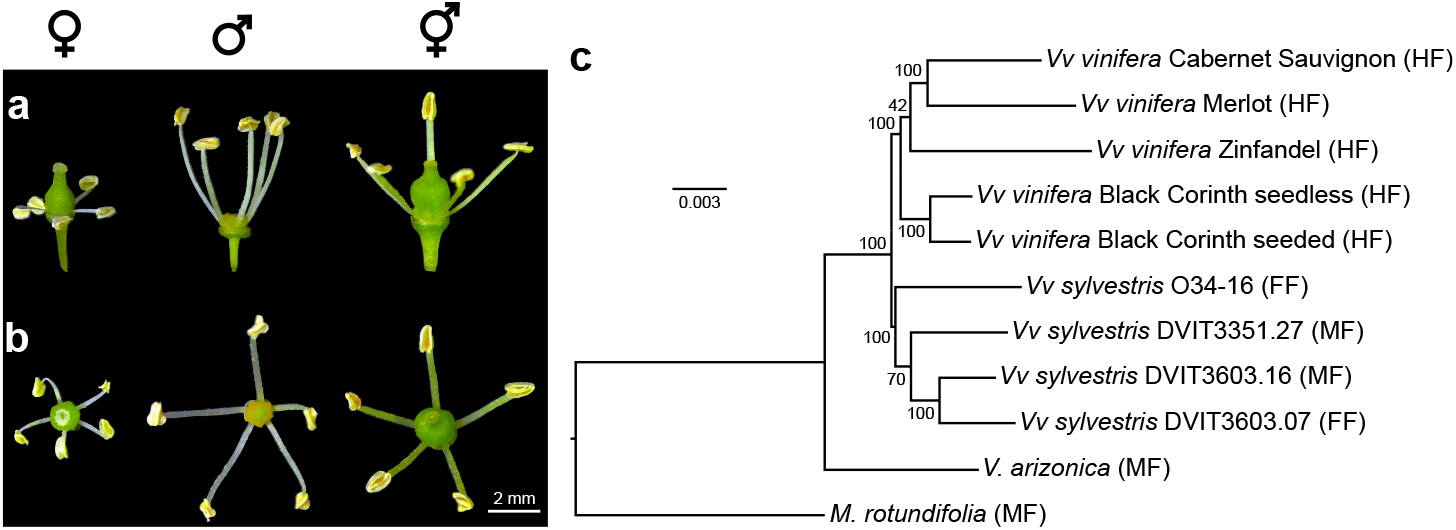
Dioecy and hermaphroditism, the morphology of flower sexes in grapevine and a phylogenetic analysis of wild and cultivated species. Side view (**a**) and top view (**b**) of the dioecious *Vv sylvestris* female O34-16 (left), male DVIT3351.27 (middle) and of the hermaphrodite *Vv vinifera* Chardonnay (right). **c**, Phylogenetic tree predicted from proteome orthology inference separates species by taxonomy and not by sex type. *M. rotundifolia* is considered an outgroup. For each genotype, allelic state of the sex locus is indicated in parenthesis.

Efforts made to understand the sex locus in grape converged on a ~150 kbp region on chromosome 2 (Dalbó *et al.*, 2000; Riaz *et al.*, 2006; Marguerit *et al.*, 2009; Fechter *et al.*, 2012; Battilana *et al.*, 2013; Picq *et al.*, 2014; Hyma *et al.*, 2015; Zhou *et al.*, 2017) and several models that explain grapevine sex inheritance have been proposed (Valleau, 1916; Oberle, 1938; Levadoux, 1946; Doazan and Rives, 1967; Antcliff, 1980; Carbonneau, 1983). In one model, males occur when a dominant allele inhibiting ovule development is present and females occur given a recessive allele associated with the inhibition of pollen development (Oberle, 1938). Another model involves a locus with three alleles that are hierarchically dominant: Male (M) > Hermaphrodite (H) > Female (F) (Valleau, 1916; Levadoux, 1946; Doazan and Rives, 1967; Antcliff, 1980; Carbonneau, 1983). This model is compatible with the previous one provided that the two genes are included in a single locus and no recombination happens between them (Zhou *et al.*, 2019a). In a third model, Carbonneau (1983) proposed a second genomic region might interact epistatically with the primary sex locus in order to explain unexpected numbers of males and females among the progeny of some hermaphrodite x hermaphrodite and female x hermaphrodite crosses.

Despite this progress, the genetic basis of sex determination in grape is elusive. The exact boundaries of the sex locus are unknown, as is/are the gene(s) involved, and whether sex determination is based on heteromorphic M, F, and H alleles. Until recently, a major limitation has been the exclusive availability of a haploid genome reference that only partially represents a female haplotype (Jaillon *et al.*, 2007; Zhou *et al.*, 2019b). Here, we report the diploid genomes of dioecious wild grapevines (males and females) and hermaphrodite cultivated grapevines. By assembling, phasing, and comparing the genomes of male, female, and hermaphrodite samples of *Vitis* spp., we characterized the sex locus structure, its gene content and variability, identified candidate fertility and sterility genes, and updated the model for grapevine sex determination.

## Results

### Phased reconstruction of the diploid sex determination locus across multiple wild and cultivated grape species

Single molecule real-time (SMRT) DNA sequencing and FALCON-Unzip (Chin *et al*., 2016) were used to assemble phased diploid genomes for three hermaphrodite *Vv vinifera* cultivars, four *Vv sylvestris* accessions (two females and two males), one male *V. arizonica*, and one male *Muscadinia rotundifolia* (**Fig. 1c**). Two additional publicly available genomes were included; the hermaphrodite *Vv vinifera* cultivars Cabernet Sauvignon and Zinfandel were sequenced with the same approach used here (Chin *et al*., 2016; Minio *et al.*, 2017, Vondras *et al*., 2019). Altogether, the dataset represented 22 haplotypes: five hermaphrodites (H), thirteen females (F), and four males (M). On average, all diploid assemblies were highly contiguous (primary assembly N50 = 1.9 ± 1.2 Mbp; haplotig N50 = 0.3 ± 0.1 Mbp) and covered a total of 931.2 ± 62.0 Mbp (*i.e.* approximatively twice the expected haploid genome size of 450 Mbp; Jaillon *et al*., 2007; **Supplementary Table 1**). Using proximity-based ligation with HiC, optical map, and multiple scaffolding methods (**Supplementary Note**), two phased copies of the nineteen chromosomes of Cabernet Sauvignon were built and used as reference for all the analysis described below. A phased, ~500 kbp sex locus was identified in each of the eleven diploid assemblies produced. This region was flanked by known markers associated with the sex locus on chromosome 2 (Riaz *et al*., 2006; Marguerit *et al*., 2009; Fechter *et al*., 2012; Picq *et al*., 2014; **Fig. 2**) and was carefully, manually annotated in each assembly to identify sex-linked polymorphisms.

**Fig. 2:**
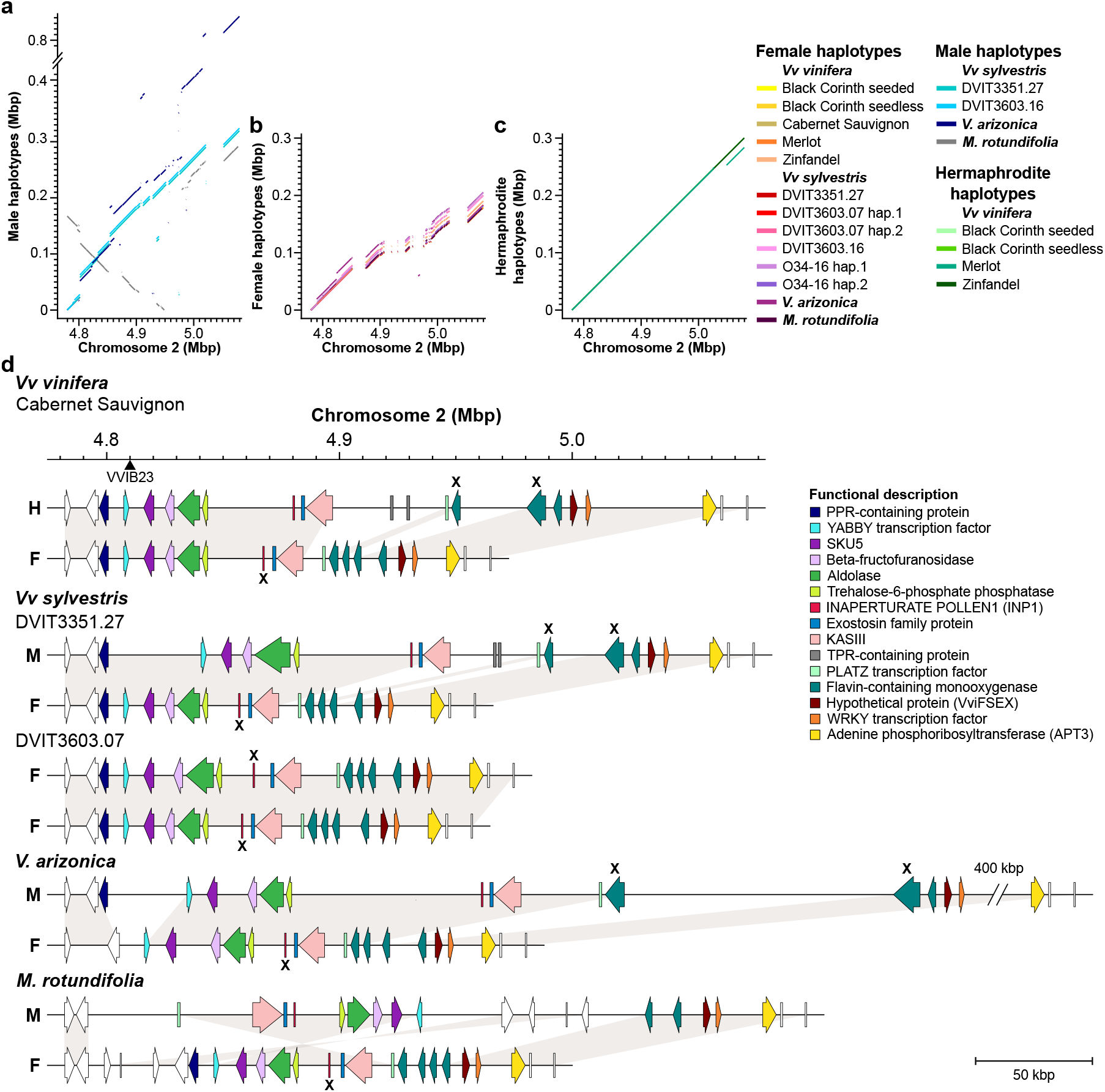
Sex-linked structural variants and their impact on gene content. Whole-sequence alignments of male (M) (**a**), female (F) (**b**), and hermaphrodite (H) (**c**) sex haplotypes on *Vv vinifera* Cabernet Sauvignon chromosome 2 Alt1 (H). **d**, From top to bottom, schematic representations of the sex locus in hermaphrodite *Vv vinifera* Cabernet Sauvignon (HF), male *Vv sylvestris* DVIT3351.27 (MF) and female *Vv sylvestris* DVIT3603.07 (FF), male *V. arizonica* (MF), and male *M. rotundifolia* (MF). A triangle along chromosome 2 marks the position of VVIB23, the genetic marker closely linked to the sex locus (Riaz *et al.*, 2006). Genes affected by nonsense mutations are indicated with an “X”.

### Sex-specific haplotype structures and gene content are conserved

Zhou *et al*. (2019b) described structural variation between Chardonnay and Cabernet Sauvignon in a portion of the sex locus. Here, with an extensive panel of phased haplotypes from wild and domesticated accessions, we successfully associate haplotype structure with sex type. The complete sex locus for both haplotypes of each accession was aligned onto Cabernet Sauvignon chromosome 2 (Alt1) and structural variants were called (SVs > 50 bp; **Fig. 2**). All sex-specific haplotypes were highly conserved throughout the dataset. Relative to the H haplotype of Cabernet Sauvignon, all F haplotypes of *Vv vinifera, Vv sylvestris, V. arizonica*, and *M. rotundifolia* shared the same four large deletions. These deletions were 22.9 kbp, 36.8 kbp, 27.7 kbp and 31.5 kbp long, respectively (**Fig. 2b**). Likewise, the structure of the M haplotype was also highly conserved in *Vv sylvestris*, *V. arizonica*, and *M. rotundifolia* grapevines (**Fig. 2a**). With the exception of a unique ~400 kbp insertion in *V. arizonica*, all M haplotypes shared two large SVs, including a 22.6 kbp insertion at position 4,802,134 and a 30 kbp deletion from positions 5,021,983 to 5,052,079. The 30 kbp deletion was also found in all F haplotypes from positions 5,021,226 to 5,052,764. Interestingly, more than half (57%) of the M sex locus was inverted in *M. rotundifolia* (**Fig. 2a**), but gene content and order within the inversion was identical to the other male haplotypes (**Fig. 2d**). Of the SVs that are shared by all haplotypes of the same sex, only two SVs altered the gene content in each of the sex haplotypes; these were (i) a deletion of two genes in F haplotypes versus H and *Vv* M haplotypes, encoding two TPR-containing proteins, and (ii) a deletion of a fourth *FLAVIN-CONTAINING MONOOXYGENASE* (*FMO*) in H and M haplotypes relative to F haplotypes. Considering that haplotypes of the same sex span multiple species and genera that diverged 47 million years ago (Zhou *et al*., 2019b), the conservation of structure and content at the locus is remarkable (**Fig. 2d**; **Extended Data Fig. 1**).

### Sex-linked polymorphisms affect protein sequences

Comparison of the *Vitis* sex haplotypes also allowed the identification of SNPs and small INDELs (≤ 50 bp) perfectly associated with sex. All sex-related polymorphisms were on chromosome 2 of Cabernet Sauvignon from positions 4,801,876 to 5,061,548 (**Fig. 3a**; **Supplementary Table 2**), further confirming and delimiting the sex locus (Fechter *et al*., 2012; Picq *et al*., 2014). In total, 1,275 SNPs were shared by all F haplotypes versus H and M haplotypes and 539 SNPs were shared by all M haplotypes versus F and H haplotypes. A small number of SNPs (127) were linked to the H haplotype. Interestingly, the highest density of M-associated SNPs was in the first 8 kbp of the sex locus (176 SNPs, 4,801,876 to 4,809,592), and the first F-associated SNP was 40 kbp downstream (4,842,196). Sex-specific SNP distributions were also similar when including *M. rotundifolia* haplotypes in the comparison, but the number of clear sex-associated SNPs decreased because of the divergence of the two genera (**Extended Data Fig. 2**). Many of the sex-linked SNPs we identified impact predicted protein sequences (**Fig. 3b**). In total, 89 non-synonymous F-specific SNPs were detected in ten genes. These included one in a gene encoding a trehalose-6-phosphate phosphatase (*TPP*), one in *INAPERTURE POLLEN1* (*INP1*), seven in a exostosin-coding gene, three in a 3-ketoacyl-acyl carrier protein synthase III gene (*KASIII*), seven in a PLATZ transcription factor-coding gene (*PLATZ*), eighteen in the first *FMO*, twenty six in the second *FMO*, eleven in the third *FMO*, eleven in the hypothetical protein (*VviFSEX*), and four in the *adenine phosphoribosyltransferase* 3 (*APT3*) (**Supplementary Table 2**). Three of these SNPs introduce a premature stop codon in the two first FMOs. Only six non-synonymous M-linked SNPs were found. There were two in the YABBY transcription factor-coding gene (*YABBY*), two in an aldolase-coding gene, one in *TPP*, and one in the third *FMO* (**Supplementary Table 2**). A phylogenetic analysis of these proteins’ sequences supports that these polymorphisms are sex-linked, with haplotypes clustering by sex rather than taxonomic proximity (**Fig. 3f**, **Extended Data Fig. 3**). All of the male YABBY and aldolase alleles formed well-supported clades separate from the F and H orthologs (**Fig. 3f**). A similar clustering pattern was observed for nine proteins that separated F from H and M. These included the TPP, INP1, the exostosin, the KASIII, the PLATZ transcription factor, the second and third FMOs, the hypothetical protein (VviFSEX), and APT3.

**Fig. 3:**
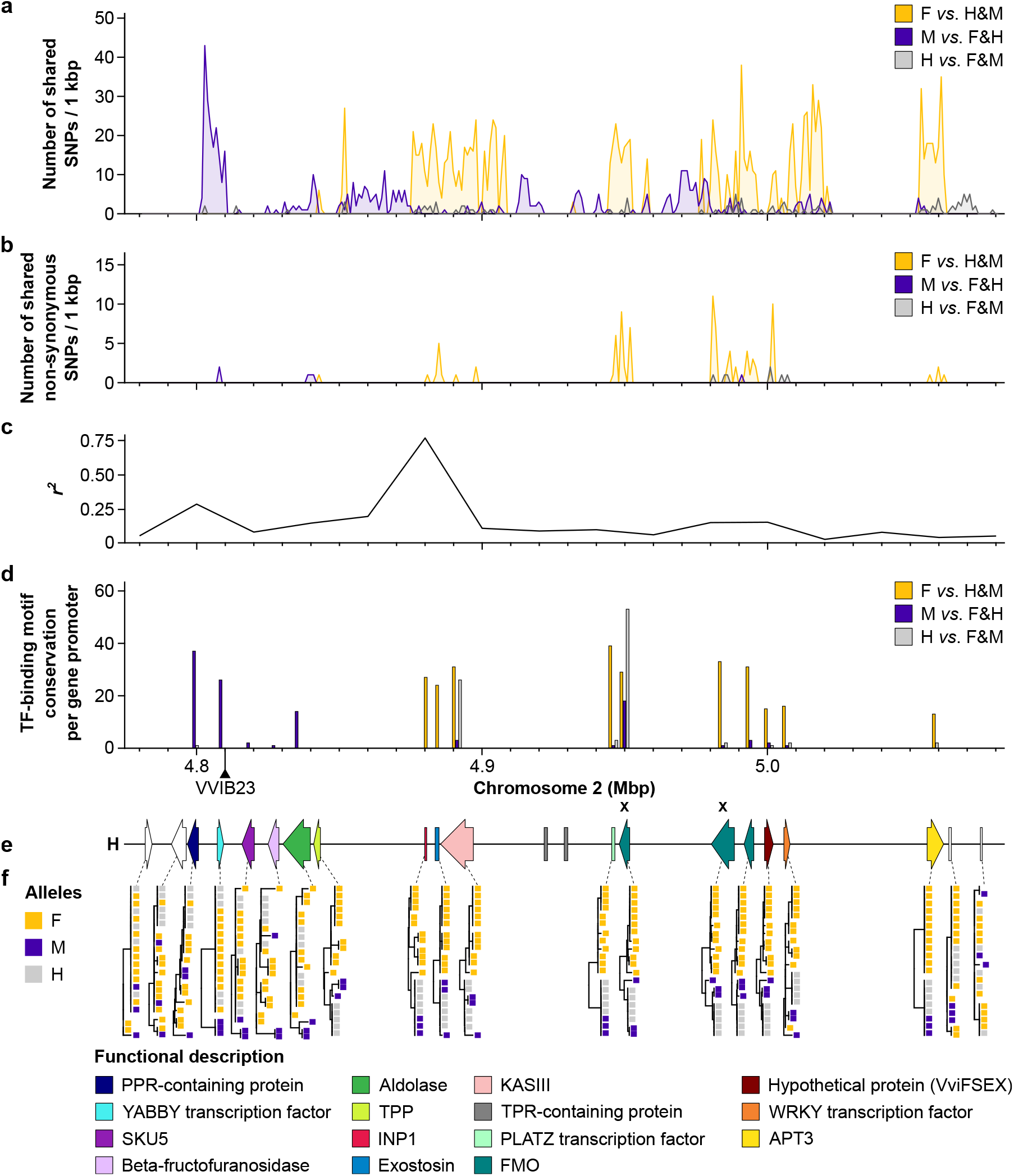
Sex-linked polymorphisms along the sex determination locus. Number of shared sex-linked SNPs per kbp (**a**) and number of non-synonymous sex-linked SNPs per kbp (**b**) detected in the sex locus. SNPs were identified aligning all *Vitis* haplotypes on *Vv vinifera* Cabernet Sauvignon Alt1 (H). **c**, Linkage disequilibrium (LD) across the locus represented as the median of the squared correlation coefficients (*r*^*2*^) between all pairs of SNPs calculated by 20 kbp windows. **d,** TF-binding motif conservation per gene promoter detected in the sex locus. Yellow, purple and grey colors are attributed to F-, M- and H-linked polymorphism respectively. A triangle along chromosome 2 marks the position of VVIB23, the genetic marker closely linked to the sex locus (Riaz *et al.*, 2006). **e**, Gene composition of the H sex locus of *Vv vinifera* Cabernet Sauvignon Alt1. **f**, Neighbor-joining trees of the protein sequences encoded by each gene of the sex determination locus. Sex haplotypes are differentiated by color. Phylogenetic trees do not include *M. rotundifolia* proteins, because they were too distant from the rest. Genes affected by nonsense mutations are indicated with an “X”.

We also detected 156 and 29 small INDELs (≤ 50 bp) shared by all F and M haplotypes, respectively. Only two of these INDELs were within exons (**Supplementary Table 3**). These were (i) a 21 bp INDEL in the first FMO-coding gene, which introduced a premature stop in H and M alleles, and (ii) an 8 bp INDEL in the grape ortholog of the *Arabidopsis thaliana INP1* (AT4G22600), which causes a frameshift and premature stop codon in all F alleles (**Fig. 4a**). A functional copy of *INP1* is necessary for fertile pollen development in *A. thaliana* (Dobritsa and Coerper, 2012). Because the same 8 bp deletion was also found in the female *M. rotundifolia* (**Fig. 4a**), this F-specific mutation likely occurred before *Vitis* and *Muscadinia* diverged and is therefore widespread in *Vitis*. PCR primers were designed to amplify markers that differentiate sex-associated alleles. *INP1* was analyzed in seven additional *Vv vinifera* (HF and HH) and two central Asian *Vitis* species, *V. piasezkii* (male; MF) and *V. romanetii* (female; FF) (**Fig. 4c**). Functional *INP1* alleles were present only in M and H haplotypes, while the non-functional *INP1* allele was only present in F haplotypes. The same markers were screened in two F1 populations. One population resulted from a cross between the female *Vv vinifera* F2-35 and the male *V. arizonica* b42-26 (186 individuals), and the other population from a cross between the female *Vv vinifera* 08326-61 and the male *Vv sylvestris* DVIT3351.27 (165 individuals). Throughout the progeny in both populations, all individuals with F flowers were homozygous for the non-functional *INP1* allele, while all M flowers carried one functional and one non-functional copy of *INP1* (**Extended Data Fig. 4,5; Supplementary Table 4**). In addition, a high linkage disequilibrium value *r*^*2*^ (0.77) was observed in this genomic region (**Fig. 3c**) across 50 *Vv vinifera* accessions (HF), suggesting a suppression of recombination at this locus. Together, these results strongly support that an 8 bp deletion in *INP1* in homozygous state causes male sterility. The nonsense mutation might be thus responsible for the absence of colpi in *Vv sylvestris* female pollen grains (Caporali *et al*., 2003; Gallardo *et al*., 2009). Abnormal pollen aperture formation has been also associated with parthenocarpy, the production of fruit without fertilization (Alva *et al*., 2015). A seeded Black Corinth cultivar and a seedless parthenocarpic Black Corinth were included in this study specifically to assess the relationship between the sex locus and parthenocarpy. Interestingly, no parthenocarpic Black Corinth-specific polymorphisms (22 SNPs and 44 INDELs) were in *INP1* or in any other coding sequences of the sex locus, indicating that sex determination and parthenocarpy are controlled separately.

**Fig. 4:**
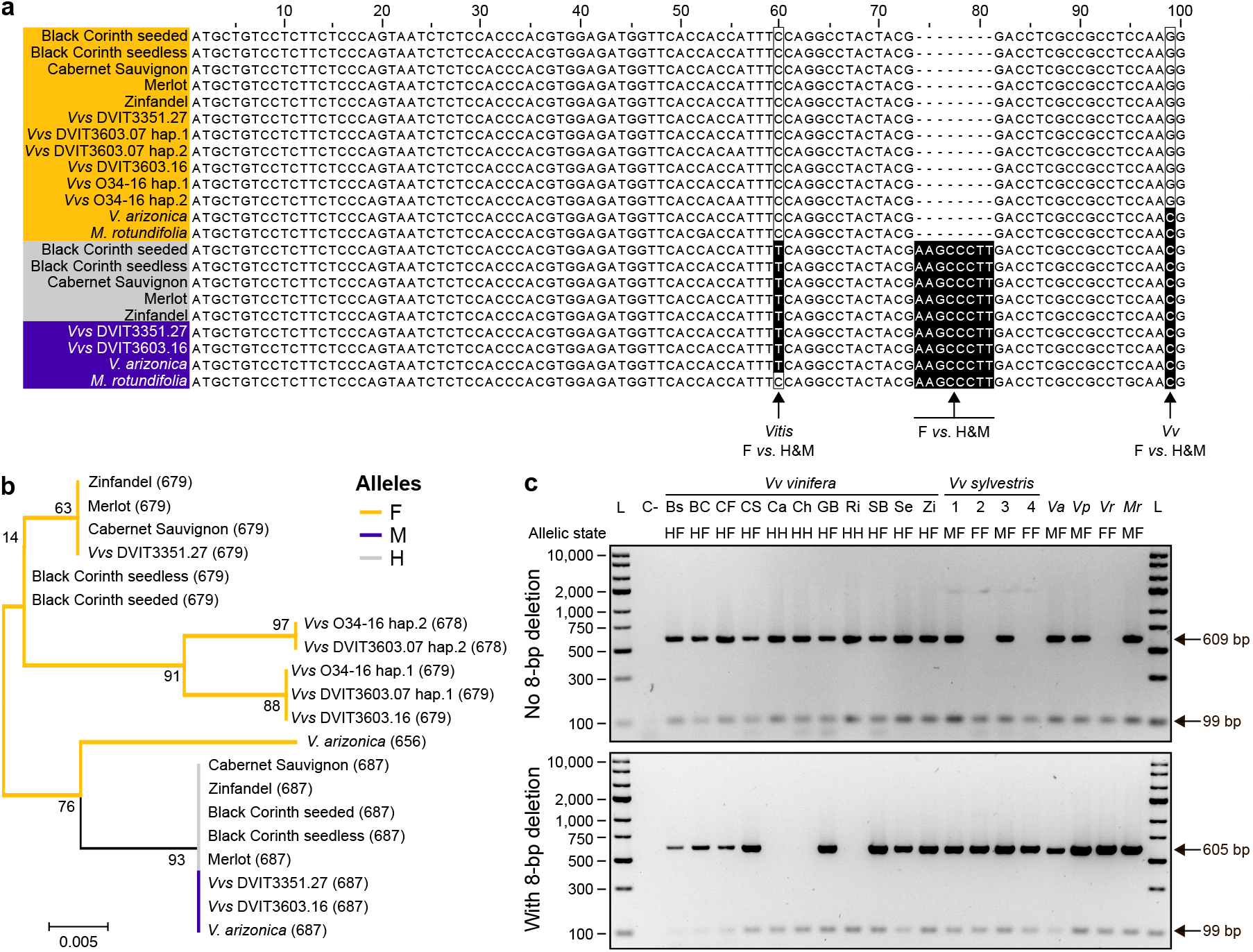
INP1, the male fertility candidate. **a**, Alignment of thirteen female, five hermaphrodite, and four male coding sequences revealed an F-linked 8 bp INDEL in *INP1*. Yellow, purple and grey colors are attributed to F, M and H haplotypes, respectively. **b**, Phylogenetic tree of the INP1 amino acid sequence. Tree branches are colored according to sex locus haplotype. Coding sequence length in bp is indicated in parentheses. **c**, Marker assay amplifying *INP1* fragment without (top panel, 609 bp) or with (bottom panel, 605 bp) the 8 bp deletion. Actin was used as a PCR positive control (99 bp fragment). Abbreviations: *Vvs*, *Vv sylvestris*; L, Ladder; C-, Negative control; *Vv vinifera*: Bs, Black Corinth seeded; BC, Black Corinth seedless; CF, Cabernet Franc; CS, Cabernet Sauvignon; Ca, Carménère; Ch, Chardonnay; GB, Gouais Blanc; Ri, Riesling; SB, Sauvignon Blanc; Se, Sémillon; Zi, Zinfandel; *Vv sylvestris*: 1, DVIT3351.27; 2, DVIT3603.07; 3, DVIT3603.16; 4, O34-16; *Va*, *V. arizonica*; *Vp*, *V. piasezkii*; *Vr*, *V. romanetii*; *Mr*, *M. rotundifolia*.

### Sex-linked polymorphisms affect transcription factor-binding sites

Sex-linked SNPs and INDELs were also found within 3 kbp region upstream of transcription start sites, including transcription factor (TF) binding sites (**Fig. 3d**; **Extended Data Fig. 6**; **Supplementary Table 5**). In total, two, nineteen, and nine TF-binding motifs were M-, F-, and H-specific, respectively. Males shared TF-binding motifs upstream of the genes encoding the PPR-containing protein, YABBY, and the aldolase that distinguished M from F and H haplotypes. Two of these motifs are associated with flowering and flower development. These included binding sites for SHORT VEGETATIVE PHASE (SVP), involved in the control of flowering time by temperature (Lee *et al*., 2007), and BES1-INTERACTING MYC-LIKE1 (BIM1), a brassinosteroid-signaling component involved in male fertility in Arabidopsis (Xing *et al*., 2013). F-linked TF-binding motifs were upstream of *INP1*, *exostosin*, *KASIII*, *PLATZ*, *FMOs*, the hypothetical protein (VviFSEX) gene, *WRKY*, and *APT3* (**Fig. 3d**; **Supplementary Table 5**).

Difference of TF-binding motif conservation in these genes’ promoters was also due to the absence of binding motifs in F haplotypes compared to H and M haplotypes. For instance, ten binding motifs corresponding to TFs involved in the control of flowering time and flower development (Schaffer *et al*., 1998; Shim *et al*., 2017) were identified only in the H and M promoter regions of *WRKY*. These included five TFs involved in the regulation of the circadian clock (AT3G09600, REVEILLE8; LHY1, LATE ELONGATED HYPOCOTYL1; EPR1, EARLY PHYTOCHROME RESPONSIVE1, also known as REVEILLE7; AT3G10113, previously shown to share a strong similarity in Arabidopsis with EPR1 (Kuno, 2003); REV1, REVEILLE1), two bHLHs (MYC2, MYC3) and three SPLs (SQUAMOSA PROMOTER BINDING PROTEIN-LIKE5, 11 and 15). Binding sites for bHLH TFs and AGAMOUS-LIKE 3 that were present in M and H haplotypes were absent from the promoter region of *APT*3 in F haplotypes, while binding sites for ARF3 (AUXIN RESPONSE FACTOR 3), involved in control of floral meristem determinacy by cytokinins (Zhang *et al*., 2018), were found only in H haplotypes (**Extended Data Fig. 7a**). These conditions suggest that genes in the *Vitis* sex locus could be differentially regulated in a sex-specific manner. In addition, the observed diversity of TF-binding motifs suggests a complex regulation of these genes involving TFs located outside the locus that could be influenced by environmental factors.

### Sex-linked genes have distinct expression patterns and are highly connected in co-expression modules

The role of genes in the sex locus during flower development was further investigated by RNA sequencing (RNA-seq) on floral buds from the hermaphrodite *Vv vinifera* Chardonnay (HH), the male and female *Vv sylvestris* DVIT3351.27 (MF) and O34-16 (FF). Flowers were sampled (i) during the early development of the reproductive structures (ii) pre-bloom during pollen maturation and (iii) at anthesis. Sequencing reads were mapped onto the phased Cabernet Sauvignon genome. Gene expression patterns during flower development were compared for sex-linked genes and alleles (**Fig. 5a**; **Supplementary Tables 6 and 7**). *INP1* was significantly more highly expressed in pre- and post-bloom female flowers compared to male and hermaphrodite flowers (adjusted P value ≤ 0.05). The high expression of *INP1* was specific to the F allele, which could reflect an attempt to compensate for the 8 bp deletion that renders the protein non-functional and causes male sterility. Conversely, *WRKY* was more highly expressed in male and hermaphrodite versus female flowers at all developmental stages. In contrast, *YABBY* was more highly expressed in male flowers compared to female and hermaphrodite flowers at the two last developmental stages. Overexpression was specific to the M allele relative to the male *sylvestris* DVIT3351.27 genome (**Supplementary Table 7**). In addition to the F-linked and M-linked sequence polymorphisms in *WRKY* and *YABBY*, respectively, expression data suggest that *WRKY* and *YABBY* are associated with male and female sterility, respectively. Curiously, *PLATZ* was more highly expressed in male and hermaphrodite flowers versus female flowers at the first developmental stage, which is consistent with the hypothesis that it could contribute to male sterility but was less expressed in male versus female and hermaphrodite flowers at the last two stages. Interestingly, *APT3* was more highly expressed in male flowers at all stages (**Extended Data Fig. 7b**). This is consistent with a previous study which showed that *APT3* overexpression was specific to carpel primordia of *sylvestris* male flowers and led the authors to propose a role in carpel abortion (Coito *et al*., 2017). The ratio of expression between H/M and F alleles was predictably very low in female flowers which only contain the F allele, was low in hermaphrodite flowers, and was high in male flowers; the expression level or “dosage” of the APT3 H/M allele might influence sex determination (**Extended Data Fig. 7c**).

**Fig. 5:**
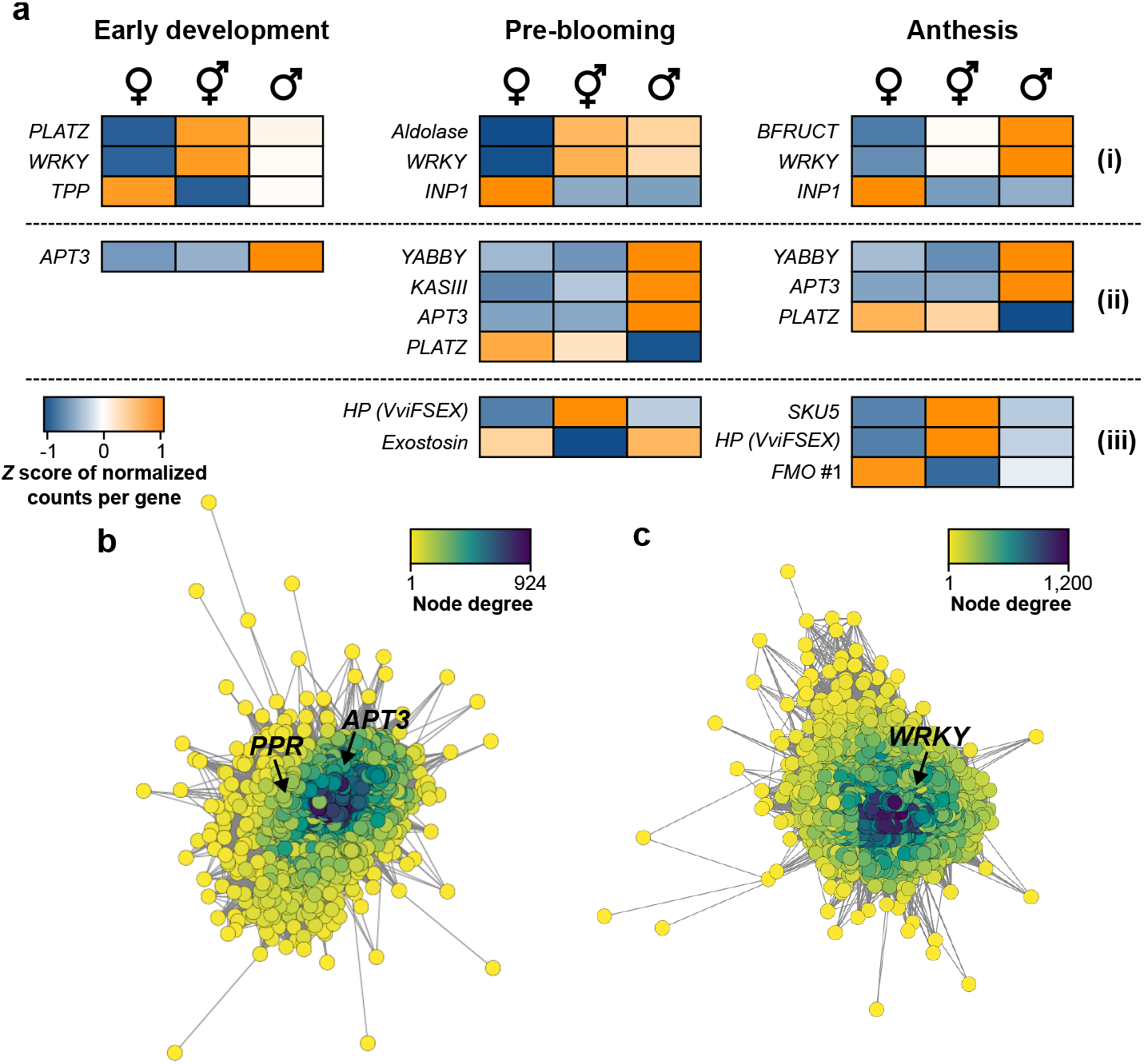
Transcriptome and gene network analyses. **a**, Sex locus genes exhibit a sex-linked expression at each floral stage. Genes are classified in three groups based on their expression pattern. Only genes differentially expressed in one flower sex compared to the two other sex types are shown. The colors of the heat map depict the *Z* score of the normalized counts per gene. Gene co-expression networks of the module ‘magenta’ **(b)**, positively correlated with M sex, and ‘red’ **(c)**, negatively correlated with F sex. Node color depicts the node degree in the network. Positions of the gene encoding a PPR-containing protein, *APT3* and *WRKY* in their corresponding networks are highlighted. Abbreviations: HP, hypothetical protein; BFRUCT, beta-fructofuranosidase.

Among the sex-linked genes are annotated transcription factors and genes associated with hormone signaling that are expected to play significant regulatory roles in co-expression networks (Serin *et al*., 2016). For this reason, Weighted Gene Co-expression Network Analysis (WGCNA; Langfelder and Horvath, 2008) was applied to assess the relationship between sex-linked genes and genes outside of the sex locus that still could play significant roles in development (**Supplementary Table 8)**. Six groups of co-expressed genes (6,830 genes in total) were positively or negatively correlated with one of the three sex haplotypes (Pearson correlation > 0.9, P value < 8e-11; **Extended Data Fig. 8**). A group of 1,176 co-expressed genes in the ‘magenta’ module (**Fig. 5b**) were well-correlated with male sex (Pearson correlation = 0.97; P value = 2e-16). This module included two genes in the sex locus encoding a PPR-containing protein and *APT3*, which shares high amino acid sequence identity with *AtAPT3* (AT4G22570). The module also included genes involved in hormonal signaling, like two uridine diphosphate glycosyltransferases (UGTs), that are orthologous to *AtUGT85A1* and *AtUGT85A3*. The F-linked *WRKY* was highly connected in a co-expression module (‘red’) of 1,511 other genes (**Fig. 5c**) that was negatively correlated with female sex (Pearson correlation = −0.92; P value = 6e-12). If WRKY plays a role in male sterility, this may occur via downregulation of the F *WRKY* allele in female flowers and associated changes in co-expression that occur in consequence. This module included an ortholog of Arabidopsis *SEPALLATA1* (*SEP1*; AT5G15800), a MADS-box gene necessary for floral organ development; SEP genes are essential for stamen development (Honma and Goto, 2000; Pelaz *et al*., 2000, 2001a, 2001b; Ditta *et al*., 2004). Both SEPALLATA1 and sex-linked WRKY are transcription factors with well-described developmental functions; this and their centrality within the ‘red’ module suggest that they may be involved in the regulation of essential components of floral development. Altogether, these data show that the expression of sex-linked polymorphisms and highly connected genes within co-expression modules participate in sex determination and other developmental processes.

## Discussion

This study constitutes an unprecedented examination of the sex determination locus in grapevine and the impact of evolution upon it. Namely, this work delineates the boundaries of and sex-specific regions of the sex determination locus, identifies conserved sex-linked polymorphisms in protein-coding genes and in their upstream regulatory regions, and updates the model of sex determination for *Vitis*. The conservation of sex-linked sequences is remarkable, and consistent with the expectation of a sex determining region largely bereft of recombination. This work was made possible by the availability of diploid genomic information and significantly expands upon knowledge concerning the *Vitis* sex locus, which was previously limited to an estimation of its approximate coordinates (Fechter *et al*., 2012; Picq *et al*., 2014), the development of sex-specific markers, and whether one or more genomic features are involved in the determination of sex (Ramos *et al*., 2014; Coito *et al*., 2017).

Overall, the model assumes that males and females observed in wild *Vitis* selections arose independently from hermaphroditic ancestors and that the hermaphroditism observed in domesticated cultivars constitutes a reversion to this sexual condition rather than a novel occurrence in the lineage (**Fig. 6**). Generally, dioecy is rare in angiosperms (Renner, 2014) and arises by either selective arrest of a single sex’s organs or by sex differentiation prior to stamen or carpel initiation (reviewed in Diggle *et al.*, 2011). Structures for both sexes are present in wild grapevine dioecious flowers, though male flowers have reduced pistils and female flowers produce infertile pollen. Earlier work on grapevine sex determination proposed that sex determination occurs after floral organ identity is established (Ramos *et al.*, 2014). Together, the existing literature supports that extant wild *Vitis* are likely derived from hermaphroditic ancestors and that dioecy results from independent and sequential male and female sterility mutations, as is predicted for angiosperms generally (Charlesworth and Charlesworth, 1978).

**Fig. 6:**
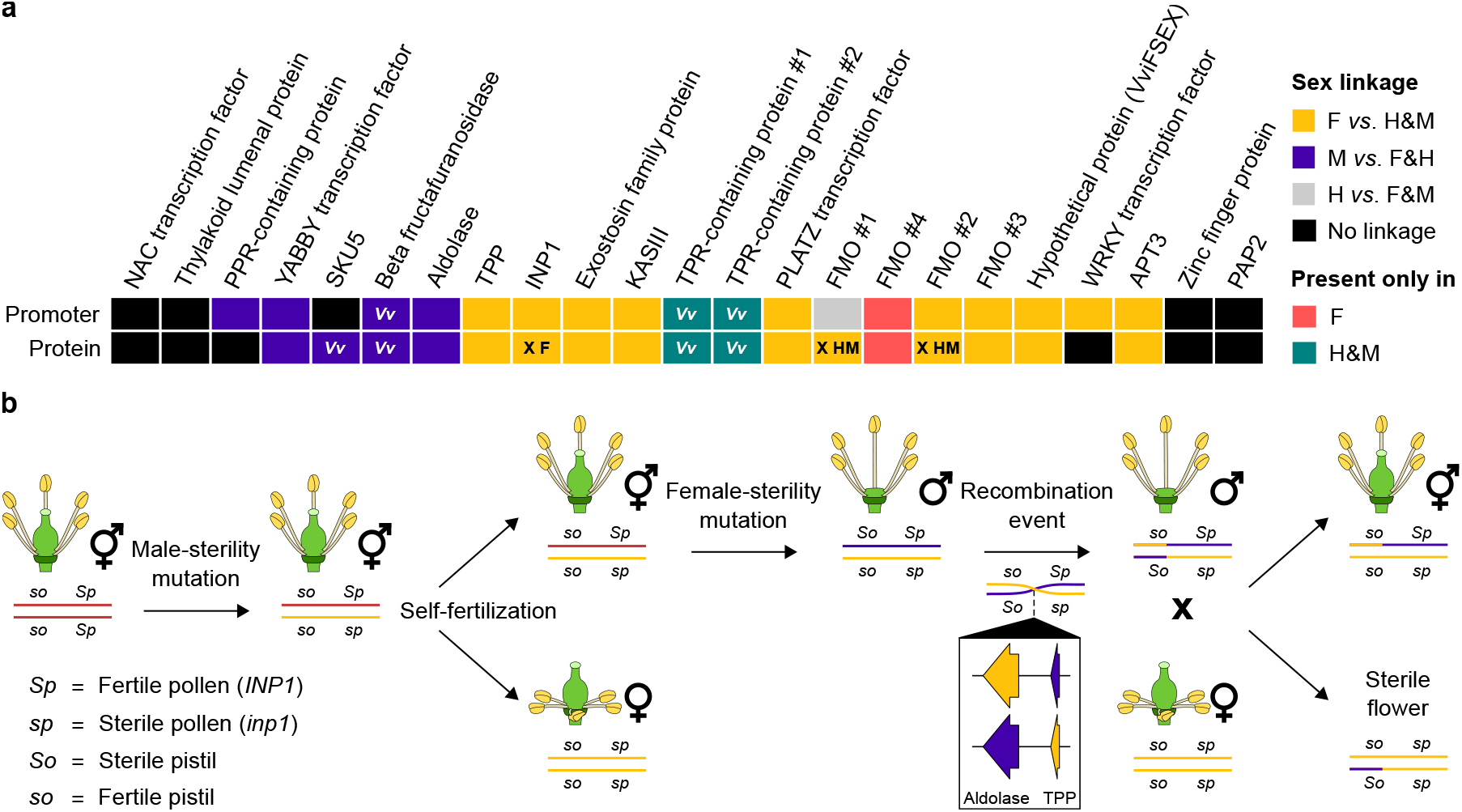
Model of the evolution of sex determination in grapevine. **a**, Graphical representation of the association between sex and the observed polymorphisms in promoter regions (top row) and encoded proteins (bottom row) of each gene present in the sex determination locus. Genes affected by nonsense mutations are indicated with an “X” and the affected haplotypes. **b**, Female dioecious flowers are thought to have evolved from a hermaphrodite ancestor through the spread of male-sterility mutation(s). Male dioecious flowers then originated as a consequence of female-sterility mutation(s). A rare recombination event was deduced to occur in the males leading to H sex determination haplotype in *Vv sylvestris*. Crossing of the aforementioned females with these males finally generate hermaphrodite flowers in domesticated *Vv vinifera* cultivars.

Within this framework, we propose two key regions in the sex locus and define their contents, each dense with genes and putative regulatory features that are male- or female-linked, and within which dominant female-sterility and recessive male-sterility mutations occurred (**Fig. 6**). The proposed mechanism of sex determination is compatible with that proposed by Oberle *et al.* (1938) involving a dominant allele inhibiting femaleness and a recessive allele inhibiting maleness. These two alleles have been designated *So* and *sp*, respectively, in keeping with the convention proposed by Oberle (1938) and reiterated by Ramos *et al.* (2014). M-linked polymorphisms occurred in regions spanning the PPR-containing protein’s promoter through an aldolase-coding gene and F-linked polymorphisms spanned *TPP* through *APT3* (**Fig. 6a**). The colocalization of these genes has not been reported previously, nor has their combined participation in sex determination to the authors’ knowledge. In each region, the sex-linked conservation of gene regulatory elements and co-expression patterns involved might contribute to the observations of Carbonneau (1983), who proposed an epistatic relationship between two loci; transcription factors could facilitate cross-talk between the two regions and widen the door for environmental signals (Golenberg and West, 2013) and stochasticity (Raj and van Oudenaarden, 2008) to influence the expression of genes carrying sex-linked polymorphisms. However, this has not been experimentally observed.

Polymorphisms in the first 8 kbp of the sex locus are predicted to include the mutation causing female sterility because these, most clearly those in and around *YABBY, SKU5*, and the aldolase-coding gene (**Fig. 3f**), distinguish M from F and H. M-linked polymorphisms were observed within or upstream of coding genes (**Fig. 3**), and YABBY exhibited an M-linked expression pattern of the M-linked allele consistent with what would be expected for a female sterility gene (**Fig. 5**). Earlier reports have associated several of these genes with pollen development and fertility. PPR is required for pollen development in rice (Liu *et al.*, 2017) and, notably, is associated with the restoration of cytoplasmic male sterility in plants that otherwise do not produce functional pollen (reviewed in Gaborieau *et al.*, 2016). *YABBY* was previously associated with sex and the VVIB23 microsatellite sequence in grapevine (Battilana *et al.*, 2013). Possibly under the regulation of unique upstream regulatory sequences, *YABBY* was more highly expressed in male flowers. Likewise, *SKU5* is up-regulated in mature cucumber pollen grains; it was proposed that SKU5 may function during cucumber floral development and pollen formation (Pawełkowicz *et al.*, 2019) and is generally associated with two-directional growth and the regulation of cell wall expansion (Sedbrook *et al.*, 2002). M-linked polymorphisms were found within an aldolase-coding gene; a proteomic study of pumpkin nectar found an aldolase uniquely in male nectar (Chatt *et al.*, 2018), and a putative aldolase- coding gene in scarlet gourd was up-regulated in response to AgNO3 treatment, which can induce stamen development in the flower buds of female plants, possibly by inhibiting ethylene response (Devani *et al.*, 2019). Based on these functions, the genes in this region appear to enhance maleness, and existing literature has not associated these genes explicitly with pistil fertility to the authors knowledge. However, this does not exclude their participation in female sterility, as polymorphisms and the expression patterns of genes in this range clearly distinguished each sex (**Fig. 3f**, **5a**).

Next, we observed a dense range of F-linked polymorphisms that distinguished F from H and M haplotypes (**Fig. 3f**) and could account for male sterility in females flowers, which includes reflexed filaments, lack of fertile pollen and pollen dimorphism (Caporali *et al*., 2003; Gallardo *et al*., 2009). Like Ramos *et al.* (2014), we propose that females arose via hermaphroditic selfing. *INP1* is involved in the formation of pollen surface apertures in Arabidopsis (Dobritsa and Coerper, 2012) and is not functional in females; this could explain the production of sterile unapertured pollen grains in females of dioecious grape species (Caporali *et al*., 2003; Gallardo *et al*., 2009). Importantly, these data show that hermaphrodite genotypes do not share the 8 bp deletion in *INP1* found in females, differing in pollen fertility, suggesting that *INP1* does not participate in pistil fertility. Moreover, all F1 males contained at least one functional *INP1* **(Fig. 4c**; **Extended Data Fig. 4, 5)**, which suggests it is required for fertile pollen.

In addition to *INP1*, our observations suggest a complex role of *APT3* and cytokinin-signaling in *Vitis* sex determination. *APT3* is highly expressed in male flowers, and sequence polymorphism shows the F allele clearly distinguished from M and H alleles. The specific overexpression of the H/M allele of *APT3* in male flowers but not in hermaphrodite flowers suggests the action of at least one secondary factor that activates *APT3* expression in males or represses its expression in hermaphrodites. The H-linked binding site for ARF3 upstream of *APT3* is a candidate. Furthermore, the distinct ratios of allelic expression (H/M:F) of this gene in each flower sex during development suggest that APT3 dosage could influence sex (**Extended Data Fig. 7c)**. APT proteins participate in cytokinin inactivation (Allen *et al*., 2002; Zhang *et al*., 2013). Cytokinins are involved in flower development and the regulation of the number of ovules per pistil, gynoecium size (Bartrina *et al*., 2011; Marsch-Martínez *et al*., 2012), and flower sex specification (Durand and Durand, 1991; Ni *et al*., 2018). In grapevine, cytokinin application restores normal female organ development, producing berries from male grapevine flowers (Negi and Olmo, 1966). Interestingly, *APT3* expression has been previously associated with the abortion of pistil structures in male grapevine flowers because of the high expression of the gene in the third and fourth whorl of male flowers (Coito *et al*., 2017). Our expression data are consistent with Coito *et al.* (2017), with uniquely high expression of *APT3* in male flowers. However, in Arabidopsis, *APT1* mutants are sterile males (Moffatt and Somerville, 1988 and Gaillard *et al*., 1998 cited in Coito *et al.*, 2017). The putative regulatory site(s) identified upstream of *APT3* are associated with hormone signaling and stamen development in Arabidopsis (**Extended Data Fig. 7a**; Ito *et al*., 2007; Song *et al*., 2013; Qi *et al*., 2015). Perhaps the polymorphisms present in the F-linked promoter of *APT3* integrate hormone signals to downregulate the gene in manner that contributes to male sterility or that is required for female fertility. However, relatively low expression of this gene in hermaphrodite flowers was not sufficient to cause male sterility. Therefore, if *APT3* participates in male sterility, then it is likely associated with the absence of expression of the H/M allele. The quantitative nature of the effect of hormones in plants suggests a role of APT3 in sex determination via the regulation of cytokinin concentrations.

The body of literature concerning the evolution of dioecy in angiosperms expects that independent male sterility and female sterility mutations (in that order) would occur in hermaphrodites. The model proposed includes this assumption, and that this would give rise to dioecy observed in extant wild *Vitis* today. Male sterility mutations would have prevailed first because this would decrease inbreeding depression and increase the quality of the progeny obtained with hermaphrodites from which they derive (Charlesworth, 2016). This pathway is more likely when selfing and inbreeding depression are relatively frequent (reviewed in Charlesworth, 2016; Käfer *et al.*, 2017). Gynodioecious species, in which females and hermaphrodites coexist, are prevalent in flowering plants (Renner, 2014). In contrast to the model proposed by Ramos *et al.* (2014) regarding the evolution of dioecy in grapevine, female-sterility mutations leading to androdioecy has significantly less support from population genetic modelling (Charlesworth and Charlesworth, 1978). Instead, high abundance of females in gynodioecious populations could facilitate the next step towards dioecy, involving sex-allocation plasticity (Delph, 2009) that would increase the “maleness” of hermaphrodites (Maurice *et al*., 1994; Ehlers and Bataillon, 2007; Spigler and Ashman, 2012; reviewed in Käfer *et al.* 2017). Furthermore, the reversion to hermaphroditism is observed for male *Vv sylvestris* flowers that bear fruit (Negi and Olmo, 1970), suggesting that gynodioecious path is more likely than the androdioecious one speculated by Ramos *et al.* (2014). For this reason, we also propose a different mechanism that accounts for the reversion to hermaphroditism in domesticates than Ramos.

Finally, we propose that the reversion of domesticated *Vv vinifera* to H arose from a rare recombination event between M- and F-linked regions in male flowers. One hypothesis regarding the rarity of dioecy among angiosperms is that dioecy easily reverts to hermaphroditism, which contrasts the notion of dioecy as a “dead end”; sterility mutations cannot be fixed because both male and female functions are necessary for sexual reproduction (reviewed in Käfer *et al.* 2017). For hermaphroditism to recur in domesticated grapevine would require recombination between the two regions. In the absence of recombination, offspring could only be male or female. Instead, recombination in males between M- and F-linked regions would make possible the hermaphrodites observed in domesticated *Vitis*, without fertile alternatives (**Fig. 6b**). Indeed, such a specific recombination is supported by these data, showing sex-linked polymorphisms are well-conserved across all sex haplotypes in dioecious wild *Vitis* and hermaphrodite domesticates.

## Materials and Methods

### Plant material

Different plant tissues were collected from several genotypes for genome sequencing, RNA sequencing and marker assay. All plant material was immediately frozen and ground to powder in liquid nitrogen after collection. For genome sequencing, young leaves were collected from three hermaphrodite *Vv vinifera* (Merlot clone FPS 15; Black Corinth with parthenocarpic fruit FPS 02.1; Black Corinth with seeded fruit, **Extended Data Fig. 9**), four dioecious *Vv sylvestris* (male DVIT3351.27 collected from Armenia; female O34-16 collected from Iran; female DVIT3603.07 and male DVIT3603.16 collected from Azerbaijan), one dioecious male *V. arizonica* (b40-14), and one dioecious male *M. rotundifolia* (‘Trayshed’). For RNA sequencing, inflorescences were collected in April and May 2019 from vines at the University of California Davis (Davis, CA, USA). These times correspond to two developmental stages (Coombe *et al.*, 1995): flowers pressed together (E-L 15) and full flowering with 50% caps off (E-L 23, **Extended Data Fig. 10**). Three genotypes were sampled, including *Vv vinifera* cv. Chardonnay clone FPS 04, male *Vv sylvestris* DVIT3351.27, and female O34-16. Floral buds were sampled from three inflorescences at E-L 15 (time 1). At E-L 23, pre- (time 2) and post-bloom (time 3) flowers were collected separately, each represented by three biological replicates. Young leaves from additional genotypes were collected for a marker assay. This included seven *Vv vinifera*, *V. piasezkii* and *V. romanetii* and two F1 populations (**Supplementary Table 9**). One F1 population was the result of a cross between the pistillate *Vv vinifera* F2-35 (Carignane x Cabernet Sauvignon) and *V. arizonica* b42-26. The other F1 population was produced by crossing female *Vv vinifera* 08326-61 (self’s of Cabernet Franc) and *Vv sylvestris* DVIT3351.27. The sexes of F1 individuals were evaluated by flower morphology and the presence of fruit.

### DNA and RNA isolation, library preparation, and sequencing

High molecular weight genomic DNA (gDNA) was isolated as in Chin *et al*. (2016) from each of the nine previously described samples. DNA purity, quantity, and integrity were evaluated with a Nanodrop 2000 spectrophotometer (Thermo Scientific, IL, USA), Qubit 2.0 Fluorometer together with the DNA High Sensitivity kit (Life Technologies, CA, USA), and by pulsed-field gel electrophoresis, respectively. SMRTbell libraries were prepared as described in Minio *et al*. (2019a) and sequenced on a PacBio Sequel system using V3 chemistry, and on a PacBio RS II (Pacific Biosciences, CA, USA) using P6-C4 chemistry for *Vv vinifera* cv. Merlot (DNA Technology Core Facility, University of California, Davis). Summary statistics of SMRT sequencing are provided in **Supplementary Table 1**. DNA extractions from the four F1 populations were performed using a modified CTAB protocol (Riaz *et al*., 2011). Total RNA were extracted from floral buds as described in Rapicavoli *et al*. (2018) from each of the three genotypes described above. cDNA libraries were prepared using the Illumina TruSeq RNA sample preparation kit v.2 (Illumina, CA, USA) and sequenced in single-end 100 bp reads on an Illumina HiSeq4000 (**Supplementary Table 10**). Four samples from the male genotype (three replicates at time 2 and one at time 3) were re-sequenced in paired-end 150 bp reads to improve library quality and depth.

### Genome assembly

Genome assembly of hermaphrodite *Vv vinifera* cv. Merlot was performed at DNAnexus (Mountain View, CA, USA) as described in Vondras *et al*. (2019; **Supplementary Note**). For each of the other eight genotypes, SMRT reads were assembled with a custom FALCON-Unzip pipeline (Minio *et al*., 2019b) reported in https://github.com/andreaminio/FalconUnzip-DClab. Repetitive regions were marked using the DAmasker TANmask and REPmask modules (Myers, 2014) before and after SMRT read error correction. Then, error-corrected reads were assembled with FALCON v.2017.06.28-18.01 (Chin *et al*., 2013). This included setting different minimum seed-read lengths (length_cutoff_pr parameter) to improve the contiguity of the primary assembly (**Supplementary Table 1**). Haplotype phasing was carried out using FALCON-Unzip with default settings. Primary contigs and haplotigs were both polished with Arrow from ConsensusCore2 v.3.0.0. To further improve sequence contiguity, primary contigs of Cabernet Sauvignon (Chin *et al*., 2016) and the nine new assemblies were scaffolded using SSPACE-Longread v.1.1 (Boetzer and Pirovano, 2014) and gaps were closed with PBJelly from PBsuite v.15.8.4 (English *et al*., 2012, 2014). Then, gene space completeness was assessed with BUSCO v.3 (Simão *et al*., 2015). Summary statistics of the ten genome assemblies is provided in **Supplementary Table 1**.

### Cabernet Sauvignon pseudomolecule construction

Additional scaffolding and gap-closing steps were performed on the Cabernet Sauvignon genome assembly. First, genome conformation was assessed using Dovetail™ Hi-C technology (Dovetail Genomics, CA, USA). A Dovetail Hi-C library was prepared as described in Lieberman-Aiden *et al*. (2009) and sequenced on an Illumina platform generating 166,370,017 150 bp paired-end reads (~100X). These data were used to scaffold the Cabernet Sauvignon assembly (after gap closing with PBJelly) with Dovetail’s HiRise™ pipeline v.1.3.0-1233267a1cde (Putnam *et al*., 2016). A diploid-aware map of Cabernet Sauvignon was generated using BioNano Genomics technology (BioNano Genomics, San Diego, CA). High molecular weight genomic DNA was extracted from Cabernet Sauvignon leaves by BioNano Genomics (San Diego, CA). DNA was then nicked and labelled using the IrysPrep Kit (BioNano Genomics, San Diego, CA). Labelled DNA was loaded onto the IrysChip nanochannel array for imaging on the Irys system (BioNano Genomics). Imaged molecules longer than 150 kbp were kept. These molecules were assembled using IrysView v.2.5.1.29842 (Shelton *et al*., 2015), generating a 1.05 Gbp consensus genome map with an N50 of 1.49 Mbp. A hybrid genome assembly was produced by combining the Cabernet Sauvignon consensus genome map with its primary contigs and haplotigs using HybridAssembler v.1.0 based on RefAligner v.5678 (Staňková *et al*., 2016). The relationship between the two haplotype assemblies was defined in micro-synteny blocks. Cross-alignments of Cabernet Sauvignon transcripts (mRNAs) were identified using GMAP v.2015-09-29 (Wu and Watanabe, 2005) and parsed with MCSCANX v.11.11.2013 (Wang *et al*., 2012) to identify collinear regions with a minimum of ten consecutive genes. Sequences lacking micro-synteny were discarded. Next, all scaffolding information obtained by the different technologies were manually combined to generate two haplotype-specific sets of mutually exclusive non-overlapping scaffolds. Finally, chromosome assignment was done based on collinearity with the *Vv vinifera* PN40024 V1 genome assembly (http://genomes.cribi.unipd.it/DATA/) using ALLMAPS v.0.7.5 (Tang *et al*., 2015) and GMAP alignments of Cabernet Sauvignon transcripts as input. The first set of pseudomolecule-scale sequences (Alt1) was the most complete and contiguous and the second set (Alt2) all of the alternative phased sequence information. Alt2 was refined by using collinearity with Alt1 to incorporate unplaced sequences; this was done using HaploSync, a tool developed in-house (https://github.com/andreaminio/HaploSync).

### Genome annotation

For all new nine assemblies, genome annotation was performed as described previously for Zinfandel (Vondras *et al*., 2019). Prior to gene prediction, RepeatMasker v.open-4.0.6 (Smit *et al*., 2013) was used with a custom *Vv vinifera* repeat library (Minio *et al*., 2019a) to identify and mask repetitive elements. Protein-coding genes were predicted using EVidenceModeler v.1.1.1 (Haas *et al*., 2008). *Ab initio* prediction was performed with several predictors (SNAP v.2006-07-28 (Korf, 2004); Augustus v.3.0.3 (Stanke *et al*., 2006); GeneMark-ES v.4.32 (Lomsadze *et al*., 2005); GlimmerHMM v.3.0.4 (Majoros *et al*., 2004); GeneID v.1.4.4 (Parra *et al*., 2000)), trained on Cabernet Sauvignon, and Augustus v.3.0.3 trained with a BUSCO dataset. Next, gene models were obtained with PASA v.2.1.0 (Haas *et al*., 2003) and by using Vitis ESTs and flcDNAs (downloaded from GenBank on 03.15.2016), *Vv vinifera* PN40024 v.V1 coding sequences (CDS; http://genomes.cribi.unipd.it/DATA/), *Vv vinifera* Tannat (TSA GAKH01.1) and *Vv vinifera* Corvina (TSA PRJNA169607) transcriptomes, and Cabernet Sauvignon corrected Iso-Seq reads (SRP132320) as transcriptional evidence.

Public RNA sequencing data were also used as transcriptional evidence for the *de novo* assembly of the Merlot, DVIT3351.27, and O34-16 transcriptomes (**Supplementary Note**). Then, EVidenceModeler v.1.1.1 (Haas *et al*., 2008) was used to integrate the *ab initio* predictions and PASA gene models with experimental evidence, which included proteins from Swissprot viridiplantae (downloaded on 2016.03.15) mapped with Exonerate v.2.2.0 (Slater and Birney, 2005) and the aforementioned transcriptional evidence aligned with GMAP v.2015-09-29 (Wu and Watanabe, 2005) and BLAT v.36×2 (Kent, 2002). Models featuring in-frame stop codons were removed. Functional annotations of the encoded proteins were assigned based on sequence homology (High Scoring Segment Pair (HSP) length > 50 amino acids) with proteins from the RefSeq plant protein database (downloaded on 2017.01.17) and domain identification by InterProScan v.5.27-66.0 (Jones *et al*., 2014). Structural annotation of the protein-coding genes in sex determination loci were manually curated. Features with fragmented functional domains were considered pseudo-genes and not subsequently considered.

### Sex determination locus localization and haplotype reconstruction

The grape sex determination locus was identified by aligning the SSR marker VVIB23 and genes previously associated with the sex determination locus (Fechter *et al*., 2012; Picq *et al*., 2014) to the chromosome-scaled Cabernet Sauvignon genome assembly. Protein-coding sequences (CDS) of Cabernet Sauvignon Alt1 in this genomic region were then aligned to the ten other genome assemblies (nine new and Zinfandel) with GMAP v.2015-09-29 (Wu and Watanabe, 2005) to identify homologous regions. When the alignments of sex locus-associated sequences were fragmented (*i.e.* with genes aligned to multiple contigs), BLAT v.36×2 (Kent, 2002) was used to determine the overlap between sequences and contigs were manually joined. Junction gaps of ten bases were added between overlaps.

A schematic representation of the haplotypes was made using the Gviz Bioconductor package v.1.20.0 (Hahne and Ivanek, 2016).

### Transcription factor-binding site analysis

For each haplotype, promoter sequences were extracted for all the genes of the sex determination locus using the R package GenomicFeatures v.1.36.4 (Lawrence *et al*., 2013) by taking a maximum of 3 kbp upstream regions from the gene transcriptional start sites (TSS). TF-binding sites were identified using the R package TFBSTools v.1.22.0 (Tan *et al*., 2016) and the JASPAR database (R package JASPAR2018 v.1.1.1; Khan *et al*., 2018).

### Linkage disequilibrium analysis

Illumina whole-genome resequencing data from 50 *vinifera* accessions from previous studies (mean depth = 21.6X; Zhou *et al*., 2017; Zhou *et al*., 2019b) were trimmed using Trimmomatic v.0.36 to remove adaptor sequences and bases for which average quality per base dropped below 20 in 4 bp window (Bolger *et al*., 2014). Trimmed paired-end reads were aligned onto Cabernet Sauvignon primary pseudo-molecules (Alt1) using Minimap2 v.2.17 (Li *et al*., 2018). Alignments with a mapping quality < 10 were removed using Samtools v.1.9 (Li *et al*., 2009). PCR duplicates introduced during library preparation were filtered in MarkDuplicates in the picard-tools v.1.119 (https://github.com/broadinstitute/picard). INDEL realignment was performed using RealignerTargetCreator and IndelRealigner (GATK v.3.5; DePristo *et al*., 2011). Sequence variants were called using HaplotypeCaller (GATK v.4.1.2.0; DePristo *et al*., 2011). SNPs were filtered using VariationFiltration (GATK v.4.1.2.0; DePristo *et al*., 2011), according to the following criteria: variant quality (QD) > 2.0, quality score (QUAL) > 40.0, mapping quality (MQ) > 30.0, and < 80% missing genotypes across all samples. Linkage disequilibrium measured as the correlation coefficient of the frequencies (*r^2^*) were calculated using Plink v.2.0 (Purcell *et al*., 2007) for pairwise SNPs with a minor allele frequency (MAF) > 0.05 across the whole sex determination region (chr2:4750000..5100000).

### Whole-sequence alignments and structural variation analysis

Pairwise alignments of all the haplotypes were performed using NUCmer from MUMmer v.4.0.0 (Marçais *et al*., 2018) and the --mum option using Cabernet Sauvignon Alt1 as a reference. Structural variants (SVs; > 50 bp) including deletions, insertions, duplications, inversions, translocations, and complex insertion-deletions (CIDs) and short INDELs (< 50 bp) were called using show-diff and show-snps, respectively. SNPs were called using show-snps from MUMmer v.4.0.0 and the −1 filter. Comparison between haplotypes for SVs and SNPs using multiinter from BEDTools v.2.19.1 (Quinlan, 2014). SNPs and INDELs were confirmed by manually inspecting the alignments of short- and long-read sequences of each genotype onto their corresponding genomes. Alignments were visualized using Integrative Genomics Viewer (IGV) v.2.4.14 and phasing the two haplotypes (Robinson *et al*., 2011).

### Phylogenetic analysis

Phylogenetic analysis of the eleven genomes was based on orthology inference. Predicted protein sequences from each genome were compared to one another using BLAST v.2.7.1+ (Altschul *et al*., 1990) and OrthoFinder v.2.3.7 (Emms and Kelly, 2015, 2019). Multiple-sequence alignments of the proteins in single-copy gene orthogroups were done with MUSCLE v.3.8.31 (Edgar, 2004). Alignments were concatenated and parsed with Gblocks v.91b, permitting up to 40 contiguous non-conserved positions and blocks no shorter than 5 amino acids (Castresana, 2000). Next, phylogenetic analysis was performed using RAxML-NG v.0.9.0 (Kozlov *et al*., 2019; Stamatakis, 2014). We applied the Maximum-Likelihood (ML) method, assumed the evolutionary model ‘LG+G8+F’, used twenty random starting trees, a boostrapping of 200 replicates and the Gblocks parsed alignments. The analysis was supervised using the Approximate-ML tree produced by OrthoFinder. Phylogenetic analysis of the proteins and promoter regions of genes in the sex locus were conducted with MEGA7 (Kumar *et al*., 2016) using the Neighbor-Joining method (Saitou and Nei, 1987) and 1,000 replicates (Felsenstein, 1985). Evolutionary distances were computed using the Poisson correction method (Zuckerkandl and Pauling, 1965) and are expressed as the number of amino acids or base substitutions per site. All positions with less than 5% site coverage were eliminated.

### Transcript abundance quantification

Illumina reads were trimmed using Trimmomatic v.0.36 (Bolger *et al*., 2014) and the following settings: LEADING:3 TRAILING:3 SLIDINGWINDOW:10:20 CROP:100 MINLEN:36. For the four samples with pair-ended reads, only the first mate was used and samples were randomly downsampled to 26 million reads with seqtk v.1.0-r57-dirty (https://github.com/lh3/seqtk) before trimming. These reads were then mapped to Cabernet Sauvignon using HISAT2 v.2.0.5 (Kim *et al*., 2015) with the following settings: --end-to-end --sensitive --no-unal -q -t --non-deterministic -k 1 (**Supplementary Table 10**). The Bioconductor package GenomicAlignments v.1.12.2 (Lawrence *et al*., 2013) was used to count reads per gene locus. Read-count normalization and statistical testing of differential expression were performed using DESeq2 v.1.16.1 (Love *et al*., 2014). Co-expression analysis was conducted with WGCNA v.1.66 (Langfelder and Horvath, 2008) using log_2_-transformed normalized counts (log_2_(normalized counts +1)). A soft-thresholding power of 12 with a scale-free model fitting index R^2^ > 0.92 was applied with a minimum module size of 30. The weighted network was converted into an unweighted network preserving all connections with topological overlap mapping metric (TOM) > 0.02, and imported into Cytoscape v.3.7.2 (Shannon *et al*., 2003).

### Marker assay

Three primers (two forward, one reverse) were designed to determine the presence or absence of an 8 bp deletion in *INP1* (INP1_HM-F, 5’-CCTACTACGAAGCCCTTGACC-3’; INP1_F-F, 5’-CAGGCCTACTACGGACCTC-3’; INP1-R, 5’-CCGTGCAGCGTTCAAATTCACTG-3’). *Actin*-specific primers (Licausi *et al*., 2010) were used to provide a positive amplification control. The PCR conditions included a 3 minute denaturation at 95 °C, followed by 35 cycles of 35 seconds at 95 °C, 30 seconds at 60 °C and 40 seconds at 72 °C, and a final extension for 5 minutes at 72 °C.

## Data availability

Sequencing data are accessible through NCBI (PRJNA593045). Genome sequences and gene annotation files are available at http://cantulab.github.io/data.

## Acknowledgments

This work was funded by the NSF grant #1741627 to B.S.G. and D.C., and partially supported by funds to DC from J. Lohr Vineyards and Wines, E. & J. Gallo Winery, the Chilean Economic Development Agency (CORFO; Project 13CEI2-21852, in collaboration with the UC Davis Chile Life Sciences Innovation Center), Viña San Pedro, Viña Concha y Toro, and the Louis P. Martini Endowment in Viticulture.

## Author contributions

M.M. and D.C. designed the project. M.M., J.L., S.R. and R.F.-B. collected samples. A.M. assembled all phased genomes. M.M., N.C., J.L., J.F.G., S.R. and R.F.-B. performed the phenotyping. R.F.-B. extracted DNA and RNA and prepared sequencing libraries. M.M. and J.L. performed the marker assay. M.D. performed optical mapping. M.M., N.C., A.M., A.M.V., J.F.G. and Y.Z. performed data analyses. M.M., N.C., A.M., A.M.V., B.S.G, and D.C. wrote the manuscript.

## References

1. Akagi, T., Henry, I. M., Tao, R. & Comai, L. A Y-chromosome–encoded small RNA acts as a sex determinant in persimmons. Science 346, 646–650 (2014).

2. Akagi, T. et al. Two Y-chromosome-encoded genes determine sex in kiwifruit. Nat. Plants 5, 801–809 (2019).

3. Allen, M., Qin, W., Moreau, F. & Moffatt, B. Adenine phosphoribosyltransferase isoforms of Arabidopsis and their potential contributions to adenine and cytokinin metabolism. Physiol Plant 115, 56–68 (2002).

4. Altschul, S. F., Gish, W., Miller, W., Myers, E. W. & Lipman, D. J. Basic local alignment search tool. J Mol Biol 215, 403–410 (1990).

5. Alva, O. et al. Pollen morphology and boron concentration in floral tissues as factors triggering natural and GA-induced parthenocarpic fruit development in grapevine. PLoS ONE 10, e0139503 (2015).

6. Antcliff, A. J. Inheritance of sex in Vitis. Annales de l’Amelioration des Plantes 30, 113–122 (1980).

7. Barrett, S. C. H. The evolution of plant sexual diversity. Nat Rev Genet 3, 274–284 (2002).

8. Battilana, J. et al. Linkage mapping and molecular diversity at the flower sex locus in wild and cultivated grapevine reveal a prominent SSR haplotype in hermaphrodite plants. Mol Biotechnol 54, 1031–1037 (2013).

9. Bartrina, I., Otto, E., Strnad, M., Werner, T. & Schmülling, T. Cytokinin regulates the activity of reproductive meristems, flower organ size, ovule formation, and thus seed yield in *Arabidopsis thaliana*. Plant Cell 23, 69–80 (2011).

10. Boetzer, M. & Pirovano, W. SSPACE-LongRead: scaffolding bacterial draft genomes using long read sequence information. BMC Bioinformatics 15, 211 (2014).

11. Bolger, A. M., Lohse, M. & Usadel, B. Trimmomatic: a flexible trimmer for Illumina sequence data. Bioinformatics 30, 2114–2120 (2014).

12. Carbonneau, A. Stérilités mâle et femelle dans le genre Vitis. I. Modélisation de leur hérédité. Agronomie 3, 635–644 (1983).

13. Caporali, E., Spada, A., Marziani, G., Failla, O. & Scienza, A. The arrest of development of abortive reproductive organs in the unisexual flower of *Vitis vinifera* ssp. *silvestris*. Sex Plant Reprod 15, 291–300 (2003).

14. Castresana, J. Selection of conserved blocks from multiple alignments for their use in phylogenetic analysis. Mol. Biol. Evol. 17, 540–552 (2000).

15. Charlesworth, B. & Charlesworth, D. A. Model for the Evolution of Dioecy and Gynodioecy. Am. Nat. 112, 975–997 (1978).

16. Charlesworth, D. Plant Sex Chromosomes. Annu Rev Plant Biol 67, 397–420 (2016).

17. Chatt, E. C. et al. Sex-dependent variation of pumpkin (*Cucurbita maxima* cv. Big Max) nectar and nectaries as determined by proteomics and metabolomics. Front. Plant Sci. 9, 860 (2018).

18. Chin, C.-S. et al. Nonhybrid, finished microbial genome assemblies from long-read SMRT sequencing data. Nat. Methods 10, 563–569 (2013).

19. Chin, C.-S. et al. Phased diploid genome assembly with single-molecule real-time sequencing. Nat. Methods 13, 1050–1054 (2016).

20. Coito, J. L. et al. *VviAPRT3* and *VviFSEX*: two genes involved in sex specification able to distinguish different flower types in *Vitis*. Front. Plant Sci. 8, 98 (2017).

21. Coombe, B. G. Growth stages of the grapevine: Adoption of a system for identifying grapevine growth stages. Aust J Grape Wine Res 1, 104–110 (1995).

22. Dalbó, M. A. et al. A gene controlling sex in grapevines placed on a molecular marker-based genetic map. Genome 43, 333–340 (2000).

23. Delph, L. F. Sex Allocation: Evolution to and from dioecy. Curr. Biol. 19, R249–R251 (2009).

24. DePristo, M. A. et al. A framework for variation discovery and genotyping using next-generation DNA sequencing data. Nat. Genet 43, 491–498 (2011).

25. Devani, R.S. et al. Flower bud proteome reveals modulation of sex-biased proteins potentially associated with sex expression and modification in dioecious *Coccinia grandis*. BMC Plant Biol. 19, 330 (2019).

26. Diggle, P. K. et al. Multiple developmental processes underlie sex differentiation in angiosperms. Trends Genet. 27, 368–76 (2011).

27. Ditta, G., Pinyopich, A., Robles, P., Pelaz, S. & Yanofsky, M.F. The SEP4 gene of *Arabidopsis thaliana* functions in floral organ and meristem identity. Curr Biol 14, 1935–1940 (2004).

28. Doazan, J. P. & Rives, M. Mise au point sur le déterminisme génétique du sexe dans le genre Vitis. Ann Amélioration Plantes 17, 105–121 (1967).

29. Dobritsa, A. A. & Coerper, D. The novel plant protein INAPERTURATE POLLEN1 marks distinct cellular domains and controls formation of apertures in the Arabidopsis pollen exine. Plant cell 24, 4452–4464 (2012).

30. Durand, B. & Durand, R. Male sterility and restored fertility in annual mercuries, relations with sex differentiation. Plant Sci 80, 107–118 (1991).

31. Edgar, R. C. MUSCLE: a multiple sequence alignment method with reduced time and space complexity. BMC Bioinformatics 5, 113 (2004).

32. Ehlers, B. K. & Bataillon, T. ‘Inconstant males’ and the maintenance of labile sex expression in subdioecious plants. New Phytol. 174, 194–211 (2007).

33. Emms, D. M. & Kelly, S. OrthoFinder: solving fundamental biases in whole genome comparisons dramatically improves orthogroup inference accuracy. Genome Biol. 16, 157 (2015).

34. Emms, D. M., & Kelly, S. OrthoFinder: phylogenetic orthology inference for comparative genomics. Genome Biol. 20, 238 (2019).

35. English, A. C. et al. Mind the Gap: Upgrading Genomes with Pacific Biosciences RS Long-Read Sequencing Technology. PLoS ONE 7, e47768 (2012).

36. English, A. C., Salerno, W. J. & Reid, J. G. PBHoney: identifying genomic variants via long-read discordance and interrupted mapping. BMC Bioinformatics 15, 180 (2014).

37. Fechter, I. et al. Candidate genes within a 143 kb region of the flower sex locus in Vitis. Mol Genet Genomics 287, 247–259 (2012).

38. Felsenstein, J. Confidence limits on phylogenies: An approach using the bootstrap. Evolution 39, 783–791 (1985).

39. Fujita, N. et al. Narrowing down the mapping of plant sex-determination regions using new Y chromosome-specific markers and heavy-ion beam irradiation induced Y deletion mutants in *Silene latifolia*. G3 2, 271–78 (2011).

40. Gaborieau, L., Brown, G. G. & Mireau, H. The propensity of Pentatricopeptide Repeat Genes to evolve into restorers of cytoplasmic male sterility. Front. Plant Sci. 7, 1816 (2016).

41. Gallardo, A., Ocete, R., Angeles López, M., Lara, M. & Rivera, D. Assessment of pollen dimorphism in populations of *Vitis vinifera* L. subsp. *sylvestris* (Gmelin) Hegi in Spain. Vitis 48, 59–62 (2009).

42. Golenberg, E. M. & West, N. W. Hormonal interactions and gene regulation can link monoecy and environmental plasticity to the evolution of dioecy in plants. Am. J. Bot. 100, 1022–1037 (2013).

43. Haas, B. J. et al. Improving the Arabidopsis genome annotation using maximal transcript alignment assemblies. Nucleic Acids Res. 31, 5654–5666 (2003).

44. Haas, B. J. et al. Automated eukaryotic gene structure annotation using EVidenceModeler and the program to assemble spliced alignments. Genome Biol. 9, R7 (2008).

45. Hahne, F. & Ivanek, R. Visualizing Genomic Data Using Gviz and Bioconductor. Methods Mol. Biol. 1418, 335–351 (2016).

46. Henry, I. M., Akagi, T., Tao, R. & Comai, L. One hundred ways to invent the sexes: theoretical and observed paths to dioecy in plants. Annu Rev Plant Biol 69, 553–575 (2018).

47. Honma, T. & Goto, K. The Arabidopsis floral homeotic gene PISTILLATA is regulated by discrete cis-elements responsive to induction and maintenance signals. Development 127, 2021–2030 (2000).

48. Hyma, K. E. et al. Heterozygous Mapping Strategy (HetMappS) for high resolution Genotyping-By-Sequencing Markers: a case study in grapevine. PLoS ONE 10, e0134880 (2015).

49. Ito, T., Ng, K.-H., Lim, T.-S., Yu, H. & Meyerowitz, E. M. The homeotic protein AGAMOUS controls late stamen development by regulating a jasmonate biosynthetic gene in Arabidopsis. Plant Cell 19, 3516–3529 (2007).

50. Jaillon, O. et al. The grapevine genome sequence suggests ancestral hexaploidization in major angiosperm phyla. Nature 449, 463–467 (2007).

51. Jones, P. et al. InterProScan 5: genome-scale protein function classification. Bioinformatics 30, 1236–1240 (2014).

52. Käfer, J., Marais, G. A. & Pannell, J. R. On the rarity of dioecy in flowering plants. Mol. Ecol. 26, 1225–1241 (2017).

53. Kent, W. J. BLAT---The BLAST-Like Alignment Tool. Genome Res. 12, 656–664 (2002).

54. Khan, A. et al. JASPAR 2018: update of the open-access database of transcription factor binding profiles and its web framework. Nucleic Acids Res. 46, D260–D266 (2018).

55. Kim, D., Langmead, B. & Salzberg, S. L. HISAT: a fast spliced aligner with low memory requirements. Nat. methods 12, 357–360 (2015).

56. Korf, I. Gene finding in novel genomes. BMC Bioinformatics 5, 59 (2004).

57. Kozlov, A. M., Darriba, D., Flouri, T., Morel, B., & Stamatakis, A. RAxML-NG: a fast, scalable and user-friendly tool for maximum likelihood phylogenetic inference. Bioinformatics 35, 4453–4455 (2019).

58. Kumar, S., Stecher, G. & Tamura, K. MEGA7: Molecular Evolutionary Genetics Analysis version 7.0 for bigger datasets. Mol. Biol. Evol. 33, 1870–1874 (2016).

59. Kuno, N. The novel MYB Protein EARLY-PHYTOCHROME-RESPONSIVE1 is a component of a slave circadian oscillator in Arabidopsis. Plant Cell 15, 2476–2488 (2003).

60. Langfelder, P. & Horvath, S. WGCNA: an R package for weighted correlation network analysis. BMC Bioinformatics 9, 559 (2008).

61. Lawrence, M. et al. Software for computing and annotating genomic ranges. PLoS Comput. Biol. 9, e1003118 (2013).

62. Lee, J. H. et al. Role of SVP in the control of flowering time by ambient temperature in Arabidopsis. Genes Dev. 21, 397–402 (2007).

63. Levadoux, L. Étude de la fleur et de la sexualité chez la vigne (C. Déhan, Montpellier, 1946).

64. Li, H. et al. The Sequence Alignment/Map format and SAMtools. Bioinformatics 25, 2078–2079 (2009).

65. Li, H. Minimap2: pairwise alignment for nucleotide sequences. Bioinformatics 34, 3094–3100 (2018).

66. Licausi, F. et al. Genomic and transcriptomic analysis of the AP2/ERF superfamily in Vitis vinifera. BMC genomics 11, 719 (2010).

67. Lieberman-Aiden, E. et al. Comprehensive mapping of long-range interactions reveals folding principles of the human genome. Science 326, 289–293 (2009).

68. Liu, Y. et al. A plastid-localized Pentatricopeptide Repeat Protein is required for both pollen development and plant growth in rice. Sci. Rep. 7, 11484 (2017).

69. Lomsadze, A., Ter-Hovhannisyan, V., Chernoff, Y. O. & Borodovsky, M. Gene identification in novel eukaryotic genomes by self-training algorithm. Nucleic Acids Res. 33, 6494–6506 (2005).

70. Love, M. I., Huber, W., Anders, S. Moderated estimation of fold change and dispersion for RNA-seq data with DESeq2. Genome Biol. 15, 550 (2014).

71. Majoros, W. H., Pertea, M. & Salzberg, S. L. TigrScan and GlimmerHMM: Two open source ab initio eukaryotic gene-finders. Bioinformatics 20, 2878–2879 (2004).

72. Maurice, S., Belhassen, E., Couvet, D. & Gouyon, P.-H. Evolution of dioecy: can nuclear–cytoplasmic interactions select for maleness? Heredity 73, 346–354 (1994).

73. Marçais, G. et al. MUMmer4: A fast and versatile genome alignment system. PLOS Comput. Biol. 14, e1005944 (2018).

74. Marguerit, E. et al. Genetic dissection of sex determinism, inflorescence morphology and downy mildew resistance in grapevine. Theor. Appl. Genet. 118, 1261–1278 (2009).

75. Marsch-Martínez, N. et al. The role of cytokinin during Arabidopsis gynoecia and fruit morphogenesis and patterning. Plant J. 72, 222–234 (2012).

76. McGovern, P. et al. Early Neolithic wine of Georgia in the South Caucasus. Proc. Natl Acad. Sci. USA 114, E10309–E10318 (2017).

77. Minio, A., Lin, J., Gaut, B. S. & Cantu D. How single molecule real-time sequencing and haplotype phasing have enabled reference grade diploid genome assembly of wine grapes. Front. Plant Sci. 8, 826 (2017).

78. Minio, A. et al. Iso-Seq allows genome-independent transcriptome profiling of grape berry development. G3 9, 755–767 (2019a).

79. Minio, A., Massonnet, M., Figueroa-Balderas, R., Castro, A. & Cantu, D. Diploid genome assembly of the wine grape Carménère. G3 9, 1331–1337 (2019b).

80. Myers, G. Efficient local alignment discovery amongst noisy long reads. In: International Workshop on Algorithms in Bioinformatics 52–67 (Springer, Berlin, 2014).

81. Negi, S. S. & Olmo, H. P. Sex conversion in a male *Vitis vinifera* L. by a kinin. Science 152, 1624–1624 (1966).

82. Negi, S. S. & Olmo, H. P. Studies on sex conversion in male *Vitis vinifera* L. (*sylvestris*). Vitis 9, 89–96 (1970).

83. Ni, J. et al. Comparative transcriptome analysis reveals the regulatory networks of cytokinin in promoting the floral feminization in the oil plant *Sapium sebiferum*. BMC Plant Biol. 18, 96 (2018).

84. Oberle, G. O. A genetic study of variations in floral morphology and function in cultivated forms of Vitis. N. Y. Agric. Exp. Sta. Tech. Bull. 250, 1–63 (1938).

85. Pannell, J. R. Plant Sex Determination. Curr. Biol. 27, R191–R197 (2017).

86. Parra, G., Blanco, E. & Guigó, R. GeneID in Drosophila. Genome Res. 10, 511–515 (2000).

87. Pawełkowicz, M. et al. Comparative transcriptome analysis reveals new molecular pathways for cucumber genes related to sex determination. Plant Reprod. 32, 193–216 (2019).

88. Pelaz, S., Ditta, G. S., Baumann, E., Wisman, E. & Yanofsky, M. F. B and C floral organ identity functions require SEPALLATA MADS-box genes. Nature 405, 200–203 (2000).

89. Pelaz, S., Gustafson-Brown, C., Kohalmi, S. E., Crosby, W. L. & Yanofsky, M. F. APETALA1 and SEPALLATA3 interact to promote flower development. Plant J. 26, 385–394 (2001a).

90. Pelaz, S., Tapia-López, R., Alvarez-Buylla, E. R. & Yanofsky, M. F. Conversion of leaves into petals in Arabidopsis. Curr. Biol. 11, 182–184 (2001b).

91. Picq, S. et al. A small XY chromosomal region explains sex determination in wild dioecious *V. vinifera* and the reversal to hermaphroditism in domesticated grapevines. BMC Plant Biol. 14, 229 (2014).

92. Pratt, C. Reproductive anatomy of cultivated grapes - A Review. Am. J. Enol. Vitic. 22, 92–109 (1971).

93. Purcell, S. et al. PLINK: a toolset for whole-genome association and population-based linkage analysis. Am. J. Hum. Genet. 81, 559–575 (2007).

94. Putnam, N. H. et al. Chromosome-scale shotgun assembly using an in vitro method for long-range linkage. Genome Res. 26, 342–350 (2016).

95. Qi, T., Huang, H., Song, S. & Xie, D. Regulation of jasmonate-mediated stamen development and seed production by a bHLH-MYB complex in Arabidopsis. Plant Cell 27, 1620–1633 (2015).

96. Quinlan, A. R. BEDTools: the Swiss-army tool for genome feature analysis. Curr. Protoc. Bioinformatics 47, 11.12.1–11.12.34 (2014).

97. Raj, A. & van Oudenaarden, A. Nature, nurture, or chance: stochastic gene expression and its consequences. Cell 135, 216–226 (2008).

98. Ramos, M. J. et al. Flower development and sex specification in wild grapevine. BMC Genomics 15, 1095 (2014).

99. Rapicavoli, J. N. et al. Lipopolysaccharide O-antigen delays plant innate immune recognition of *Xylella fastidiosa*. Nat. Commun. 9, 390 (2018).

100. Renner, S. S. The relative and absolute frequencies of angiosperm sexual systems: Dioecy, monoecy, gynodioecy, and an updated online database. Am. J. Bot. 101, 1588–1596 (2014).

101. Renner, S. S. Pathways for making unisexual flowers and unisexual plants: Moving beyond the “two mutations linked on one chromosome” model. Am. J. Bot. 103, 587–589 (2016).

102. Riaz, S., Krivanek, A. F., Xu, K. & Walker, M. A. Refined mapping of the Pierce’s disease resistance locus, PdR1, and Sex on an extended genetic map of *Vitis rupestris* × *V. arizonica*. Theor. Appl. Genet. 113, 1317–1329 (2006).

103. Riaz, S., Tenscher, A. C., Ramming, D. W. & Walker, M. A. Using a limited mapping strategy to identify major QTLs for resistance to grapevine powdery mildew (*Erysiphe necator*) and their use in marker-assisted breeding. Theor. Appl. Genet. 122, 1059–1073 (2011).

104. Robinson, J. T. et al. Integrative genomics viewer. Nat. Biotechnol. 29, 24–26 (2011).

105. Saitou, N. & Nei, M. The neighbor-joining method: A new method for reconstructing phylogenetic trees. Mol. Biol. Evol, 4, 406–425 (1987).

106. Schaffer, R. et al. The late elongated hypocotyl mutation of Arabidopsis disrupts circadian rhythms and the photoperiodic control of flowering. Cell 93, 1219–1229 (1998).

107. Sedbrook, J. C., Carroll, K. L., Hung, K. F., Masson, P. H., & Somerville, C. R. The Arabidopsis SKU5 gene encodes an extracellular glycosyl phosphatidylinositol-anchored glycoprotein involved in directional root growth. Plant cell 14, 1635–1648 (2002).

108. Serin, E. A., Nijveen, H., Hilhorst, H. W. & Ligterink, W. Learning from co-expression networks: possibilities and challenges. Front. Plant Sci. 7, 444 (2016).

109. Simão, F. A., Waterhouse, R. M., Ioannidis, P., Kriventseva, E. V. & Zdobnov, E. M. BUSCO: assessing genome assembly and annotation completeness with single-copy orthologs. Bioinformatics 31, 3210–3212 (2015).

110. Shannon, P. et al. Cytoscape: a software environment for integrated models of biomolecular interaction networks. Genome Res. 13, 2498–2504 (2003).

111. Shelton, J. M. et al. Tools and pipelines for BioNano data: Molecule assembly pipeline and FASTA super scaffolding tool. BMC Genomics 16, 1–16 (2015).

112. Shim, J. S., Kubota, A. & Imaizumi, T. Circadian clock and photoperiodic flowering in Arabidopsis: CONSTANS is a hub for signal integration. Plant Physiol. 173, 5–15 (2017).

113. Slater, G. S. C. & Birney, E. Automated generation of heuristics for biological sequence comparison. BMC Bioinformatics 6, 31 (2005).

114. Smit, A. F. A., Hubley, R. & Green, P. RepeatMasker Open-4.0. http://www.repeatmasker.org 2013–2015 (2013).

115. Song, S., Qi, T., Huang, H. & Xie, D. Regulation of stamen development by coordinated actions of jasmonate, auxin, and gibberellin in Arabidopsis. Mol. Plant 6, 1065–1073 (2013).

116. Spigler, R. B., Lewers, K. S., Main, D. S. & Ashman, T. L. Genetic mapping of sex determination in a wild strawberry, *Fragaria virginiana*, reveals earliest form of sex chromosome. Heredity 101, 507–517 (2008).

117. Spigler, R. B. & Ashman, T.-L. Gynodioecy to dioecy: are we there yet? Ann. Bot. 109, 531–543 (2012).

118. Stamatakis, A. RAxML version 8: a tool for phylogenetic analysis and post-analysis of large phylogenies. Bioinformatics 30, 1312–1313 (2014).

119. Stanke, M. et al. AUGUSTUS: ab initio prediction of alternative transcripts. Nucleic Acids Res. 34, W435–W439 (2006).

120. Staňková, H. et al. BioNano genome mapping of individual chromosomes supports physical mapping and sequence assembly in complex plant genomes. Plant Biotechnol. J. 14, 1523–1531 (2016).

121. Tan, G. & Lenhard, B. TFBSTools: an R/bioconductor package for transcription factor binding site analysis. Bioinformatics 32, 1555–1556 (2016).

122. Tang, H. et al. ALLMAPS: Robust scaffold ordering based on multiple maps. Genome Biol. 16, 1–15 (2015).

123. This, P., Lacombe, T. & Thomas, M. R. Historical origins and genetic diversity of wine grapes. Trends Genet. 22, 511–519 (2006).

124. Valleau, W. D. Inheritance of sex in the grape. Am. Nat. 50, 554–564 (1916).

125. Vondras, A. M. et al. The genomic diversification of clonally propagated grapevines. Preprint at https://www.biorxiv.org/content/10.1101/585869v1 (2019).

126. Wang, Y. et al. MCScanX: a toolkit for detection and evolutionary analysis of gene synteny and collinearity. Nucleic Acids Res. 40, 1–14 (2012).

127. Westergaard, M. The mechanism of sex determination in dioecious plants. Adv. Genet. 9, 217–281 (1958).

128. Wu, T. D. & Watanabe, C. K. GMAP: a genomic mapping and alignment program for mRNA and EST sequences. Bioinformatics 21, 1859–1875 (2005).

129. Xing, S. et al. SPL8 acts together with the brassinosteroid-signaling component BIM1 in controlling *Arabidopsis thaliana* male fertility. Plants (Basel) 2, 416–428 (2013).

130. Zhang, K. et al. AUXIN RESPONSE FACTOR3 regulates floral meristem determinacy by repressing cytokinin biosynthesis and signaling. Plant Cell 30, 324–346 (2018).

131. Zhang, X. et al. Adenine phosphoribosyl transferase 1 is a key enzyme catalyzing cytokinin conversion from nucleobases to nucleotides in Arabidopsis. Mol. Plant 6, 1661–1672 (2013).

132. Zhou, Y., Massonnet, M., Sanjak, J. S., Cantu, D. & Gaut, B. S. Evolutionary genomics of grape (*Vitis vinifera* ssp. *vinifera*) domestication. Proc. Natl Acad. Sci. USA 114, 11715–11720 (2017).

133. Zhou, Y., Muyle, A. & Gaut, B. S. Evolutionary Genomics and the Domestication of Grapes. In The Grape Genome 39–55 (Spinger, Cham, 2019a).

134. Zhou, Y. et al. The population genetics of structural variants in grapevine domestication. Nat. Plants 5, 965–979 (2019b).

135. Zuckerkandl, E. & Pauling, L. Evolutionary divergence and convergence in proteins. In Evolving Genes and Proteins 97–166 (Academic Press, New York, 1965).

